# Transcriptomic, protein-DNA interaction, and metabolomic studies of VosA, VelB, and WetA in *Aspergillus nidulans* asexual spores

**DOI:** 10.1101/2020.09.09.290809

**Authors:** Ming-Yueh Wu, Matthew E. Mead, Mi-Kyung Lee, George F. Neuhaus, Donovon A. Adpressa, Julia I. Martien, Ye-Eun Son, Heungyun Moon, Daniel Amador-Noguez, Kap-Hoon Han, Antonis Rokas, Sandra Loesgen, Jae-Hyuk Yu, Hee-Soo Park

## Abstract

In filamentous fungi, asexual development involves morphological differentiation and metabolic changes leading to the formation of asexual spores. The process of asexual spore formation in *Aspergillus* is precisely regulated by multiple transcription factors (TFs), including VosA, VelB, and WetA, and these three TFs are key regulators of the formation and maturation of asexual spores (conidia) in *Aspergillus* including the model fungus *Aspergillus nidulans*. To gain a mechanistic insight on the complex regulatory roles of these TFs in asexual spores, we conducted genome-wide studies on the expression, protein-DNA interactions, and primary and secondary metabolism employing *A. nidulans* conidia. RNA sequencing and chromatin immunoprecipitation-sequencing data have revealed that the three TFs directly or indirectly regulate the expression of genes associated with spore-wall formation/integrity, asexual development, and secondary metabolism. In addition, metabolomics analyses of wild-type and mutant conidia indicate that these three TFs regulate a diverse array of primary and secondary metabolism. In summary, WetA, VosA, and VelB play inter-dependent and distinct roles governing morphological development and primary/secondary metabolic remodeling in *Aspergillus* conidia.

**Importance:** Filamentous fungi produce a vast number of asexual spores that act as reproductive and propagator cells. These spores affect humans, due to the infectious or allergenic nature of the propagule. *Aspergillus* species produce asexual spores called conidia and their formation involves morphological development and metabolic changes, and the associated regulatory systems are coordinated by spore-specific transcription factors. To understand the underlying global regulatory programs and cellular outcomes associated with conidia formation, functional genomic and metabolomic analyses were performed in the model fungus *Aspergillus nidulans*. Our results show that the fungus specific WetA/VosA/VelB transcription factors govern the coordination of morphological and chemical developments during sporogenesis. The results of this study provide insights into the genetic regulatory networks about how morphological developments and metabolic changes are coordinated in fungi. The findings are relevant for other *Aspergillus* species such as the major human pathogen *Aspergillus fumigatus* and the aflatoxin-producer *Aspergillus flavus*.

## Introduction

Fungal asexual spores (conidia) are key reproductive cells that are essential to the long-term survival of filamentous fungi under a variety of environmental conditions (1). Conidia can easily disperse into various environmental niches and act as infectious units for some pathogenic fungi (2-4). Asexual spore formation (conidiogenesis) involves developmental and metabolic changes in the organism and the associated regulatory systems are precisely coordinated (5, 6). Current knowledge about conidiogenesis is derived from numerous studies of model filamentous fungi such as *Aspergillus nidulans* (7-10).

The entire process of conidiogenesis is regulated by distinct gene sets, including central, upstream, and feedback regulators (6, 11). These components are highly conserved in *Aspergillus* species (12). In order to initiate conidiation, upstream developmental activators (FluG and FlbA-E) induce mRNA expression of *brlA*, a key initiator of conidiogenesis (13); whereas, several negative regulators such as SfgA and NsdD repress when the fungal hyphae acquire developmental competence (14-16). After the initiation of conidiogenesis, BrlA, AbaA, and WetA sequentially control the conidiation-specific genetic regulatory network, thereby regulating the formation of asexual specialized structures called conidiophores, which consist of aerial stalks, vesicles, metulae, phialides, and conidia (9, 17). These regulators are considered to form the central regulatory pathway (BrlA→AbaA→WetA) in *Aspergillus* species (18). BrlA is a key transcription factor which activates the expression of *abaA* and other genes in the early stage of conidiation (19, 20). AbaA is a TEF1 (transcriptional enhancer factor-1) family transcription factor governing the expression of certain genes such as *wetA, vosA, velB*, and *rodA* in the metulae and phialides (21-23). WetA plays an important role in conidial wall integrity and conidial maturation during the late phase of conidiogenesis (24, 25). Our recent studies have shown that WetA functions as a DNA-binding protein that regulates spore-specific gene expression (25, 26). Along with WetA, two velvet regulators, VosA and VelB, which are fungus-specific transcription factors, coordinate morphological, structural, and chemical developments, as well as exert feedback control of BrlA in conidia (27-30).

Previous studies have found that single knockout mutants of *vosA, velB*, and *wetA* share multiple conidial phenotypes including reduced spore viability, impaired trehalose biosynthesis, defective cell wall integrity, and reduced stress tolerance (25, 31, 32). Results of chromatin immunoprecipitation analyses have demonstrated that VosA and WetA recognize certain DNA-sequences(s) in the promoter regions of target genes and regulate mRNA expression of spore-specific genes in asexual spores (25, 29). In addition, the deletion of *vosA* or *wetA* affects transcript expression in secondary metabolite cluster genes (25, 30, 33). Biochemical studies have determined that VosA interacts with VelB in conidia, and this complex controls trehalose and β-glucan biosynthesis (30, 34). Importantly, the roles of these three transcription factors are conserved in *Aspergillus* species (35-38). Considered jointly, these results suggest that VosA, VelB, and WetA are key transcription factors that orchestrate spore-specific gene expression in *A. nidulans*. Although the role of each regulator has been studied, the regulatory networks between these proteins have not, to date, been investigated in detail. In addition, the effects of these three proteins on primary and secondary metabolism are yet to be elucidated.

In this study, we aimed to determine the cross-regulatory mechanisms of VosA/VelB/WetA in fungal conidiation using comparative transcriptomic and metabolomic analyses of the wild-type (WT) and null mutants of *wetA, velB*, and vosA in *A. nidulans* conidia. In addition, the direct targets of these regulators were identified by combining the results from the VosA- and VelB-chromatin interactions using chromatin immunoprecipitation sequencing (ChIP-seq) analysis with WetA-direct targets identified in a previous study (25). The results clarify the detailed molecular mechanisms by which VosA/VelB and WetA control defined common and distinct regulons and increase the overall understanding of the regulatory networks that govern fungal cell differentiation and metabolism.

## Results

### VosA-, VelB-, and WetA-mediated gene regulation in *A. nidulans* conidia

To understand the conserved and divergent regulatory roles of VosA, VelB, and WetA *in A. nidulans* conidia, comparative analysis of gene expression differences between the WT and null mutant conidia was carried out **(Figure 1)**. The 40.98% (4503/10988), 45.61% 5012/10988), and 51.96% (5729/10988) genes of the *A. nidulans* genome are differentially regulated in the Δ*vosA*, Δ*velB*, and Δ*wetA* mutant conidia, respectively, suggesting that the three regulators have a broad regulatory effect in conidia **(Figure S1)**. A total of 2143 differentially expressed genes (DEGs) between the WT and the Δ*vosA*, Δ*velB*, and Δ*wetA* mutant conidia were identified (**Figure 1A**, fold change > 2.0 for upregulation or downregulation, and q-value < 0.05). The mRNA expression levels of 890 genes were downregulated in all three mutant conidia, compared with the WT conidia. However, in all three mutant conidia, the mRNA levels of 1253 genes were upregulated. Among them, mRNA expression of a variety of genes associated with asexual development and signal transduction was affected by these three transcription factors **(Tables S1-S2)**. Importantly, 748 and 769 DEGs were down- or up-regulated by both Δ*vosA* and Δ*velB* mutant conidia, but not Δ*wetA* mutant conidia, respectively, while the mRNA levels of 2792 genes were affected solely in the *wetA* null mutant conidia. Put together, these results suggest that VosA and VelB share more DEGs while the WetA regulon has many more uniquely regulated genes.

**Figure 1.**
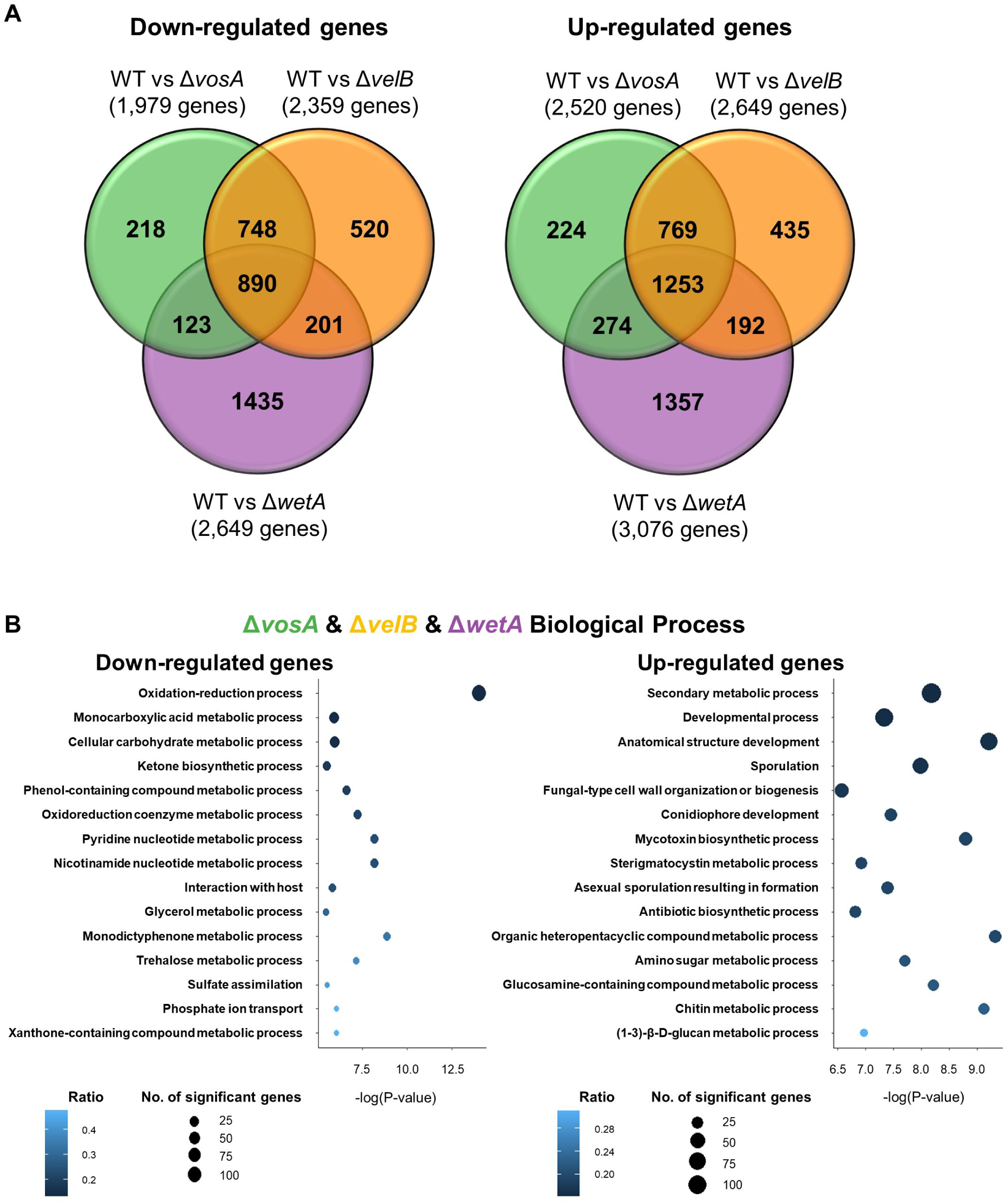
Genome-wide analyses of the genes differentially affected by VosA, VelB, and WetA in *A. nidulans* conidia. **(A)** Venn diagram showing the genes whose mRNA levels are down-regulated (left) or up-regulated (right) by the absence of VosA, VelB, or WetA in conidia. **(B)** Gene Ontology (GO) term enrichment analysis of down-regulated (left) or up-regulated (right) genes the Δ*vosA*, Δ*velB*, and Δ*wetA* conidia.

To further gain insight into the regulatory roles of these transcription factors, functional category analyses using Gene Ontology (GO) terms were carried out **(Figure 1B)**. Results of the GO analysis demonstrated that several genes involved in the monocarboxylic acid metabolic process, oxidation-reduction process, trehalose metabolic process, and cellular carbohydrate metabolic process were downregulated in all three mutant conidia, whereas a large number of genes associated with the secondary metabolic biosynthetic process, chitin biosynthetic process, asexual sporulation resulting in formation, and (1-3)-β-D-glucan metabolic process were upregulated in these mutant conidia. The VosA- and VelB-specific downregulated genes were enriched in functional categories that included cellular catabolic process, protein localization, and acetate catabolic process. The functional GO categories associated with the VosA- and VelB-specific upregulated genes were the secondary metabolic biosynthetic process, steroid metabolic process, and transport **(Figure S2A)**. Interestingly, a large number of genes involved in the RNA metabolic process was downregulated in Δ*wetA* mutant conidia but not in Δ*vosA* or Δ*velB* mutant conidia **(Figure S2B)**.

### Putative direct targets of VosA, VelB, and/or WetA in conidia

Our previous studies reported that VosA contains the velvet DNA binding domain, which recognizes the VosA binding motif in certain promoter regions (29). To identify the VelB direct target genes and compare the putative direct target genes of VosA and VelB, ChIP experiments, followed by high-throughput sequencing of the enriched DNA fragments were carried out. ChIPs from strains containing FLAG epitope-tagged versions of VosA and VelB were compared to ChIPs from wild type conidia that did not contain the FLAG epitope. A total of 1,734 and 655 genes that were VosA- and VelB-peak associated, respectively, were identified using the same analysis pipeline as described previously (25) **(Figure 2)**. To identify the VosA/VelB response elements, DNA sequences in the 100 bp surrounding each peak were subjected to Multiple Em for Motif Elicitation (MEME) analysis, which led to the predicted VosA-response-element (VoRE) and the predicted VelB-response-element (VbRE) **(Figure 2A)**. Interestingly, the predicted VbRE (5’-CCXTGG-3’) was quite similar to the predicted VoRE (5’-CCXXGG-3’).

**Figure 2.**
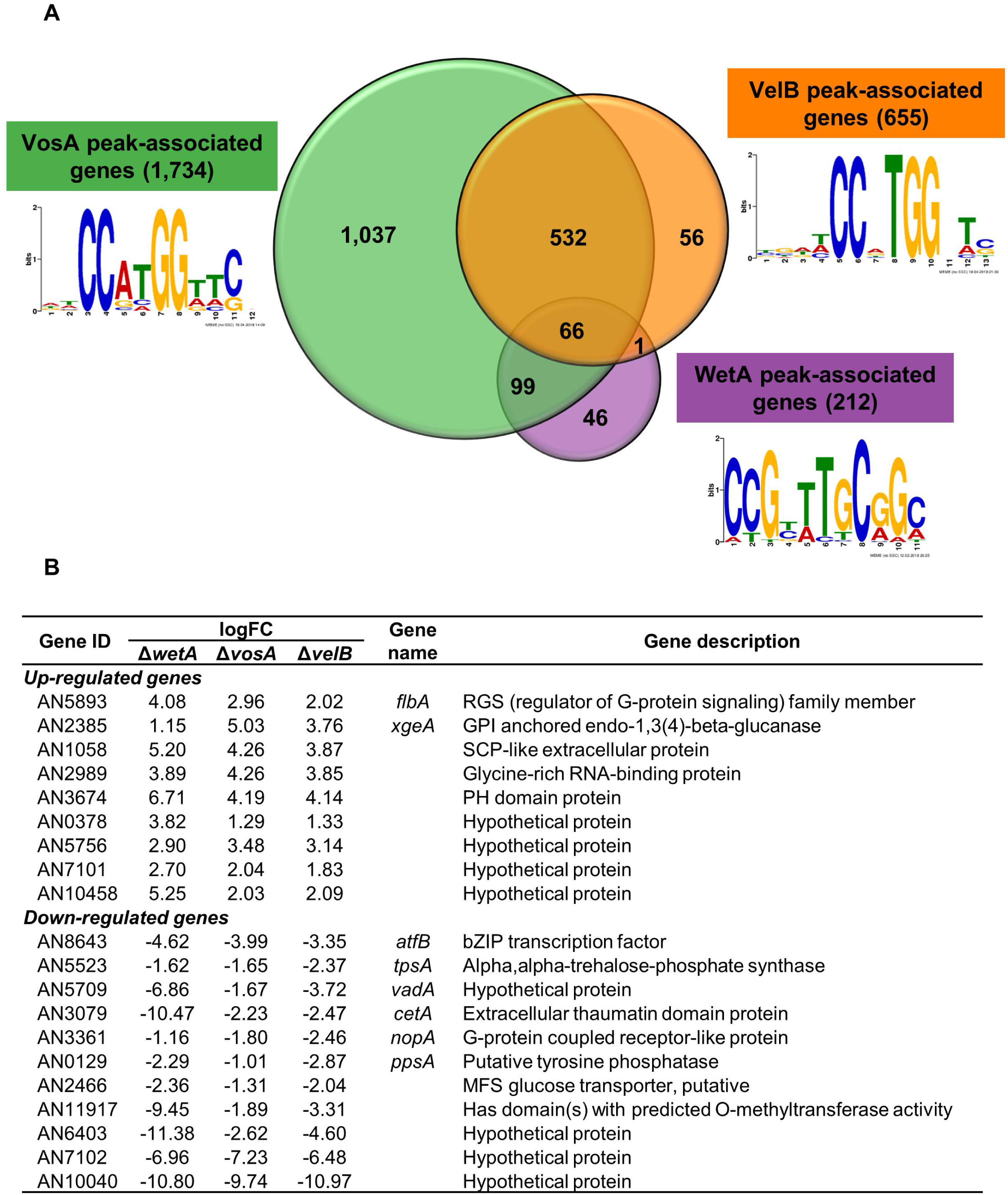
Identification of VosA, VelB, and WetA direct targets in *A. nidulans* conidia. **(A)** Venn diagram showing the number of the VosA, VelB, and WetA peak-associated genes in conidia. Motifs identified in peak-associated genes are shown next to the labels. (**B**) Summary of potential VosA, VelB, and WetA direct target DEGs in *A. nidulans* conidia.

We then compared the results of the ChIP-seq and RNA-seq analyses to identify potential direct target genes of the three transcription factors **(Table S3)**. There were 66 genes associated with the peaks of all three transcription factors **(Table S4)**. Among them, 22 genes, including *flbA, xgeA, atfB, tpsA, vadA, cetA, nopA*, and *ppsA*, were DEGs in all three null mutants **(Figure 2B)**. Importantly, 532 genes were considered to be potential direct target genes for both VosA and VelB but not WetA. A total of 166 genes were upregulated in both Δ*vosA* and Δ*velB* mutant conidia. These genes, including *brlA, fadA, rosA, steA, steC*, and *veA*, were found primarily to be involved in asexual or sexual developmental processes. Taking these results together with the previous results (27, 34), we suggest that VosA works with VelB and the VosA-VelB complex coordinates the processes involved in conidial production and maturation in *A. nidulans*.

### The roles of VosA, VelB, and/or WetA in conidial wall integrity

Previous studies have shown that the deletion of *vosA, velB*, or *wetA* alters the amount of trehalose and β-glucan in conidia (25, 30), suggesting that these genes play a conserved role in regulating the mRNA expression of genes associated with conidial integrity. High-performance liquid chromatography (HPLC) analysis demonstrated that the trehalose contents of the three null mutant conidia were dramatically decreased **(Figure 3A)**. In addition, the mRNA expression of most genes involved in trehalose biosynthesis was downregulated **(Figure 3B & Table S5)**. Moreover, most genes associated with chitin and β-(1,3)-glucan biosynthesis were upregulated in the Δ*vosA*, Δ*velB*, and Δ*wetA* mutant conidia **(Figure 3C and 3D)**. These results suggest that VosA, VelB, and WetA govern the mRNA expression of genes associated with conidial wall integrity in *A. nidulans*.

**Figure 3.**
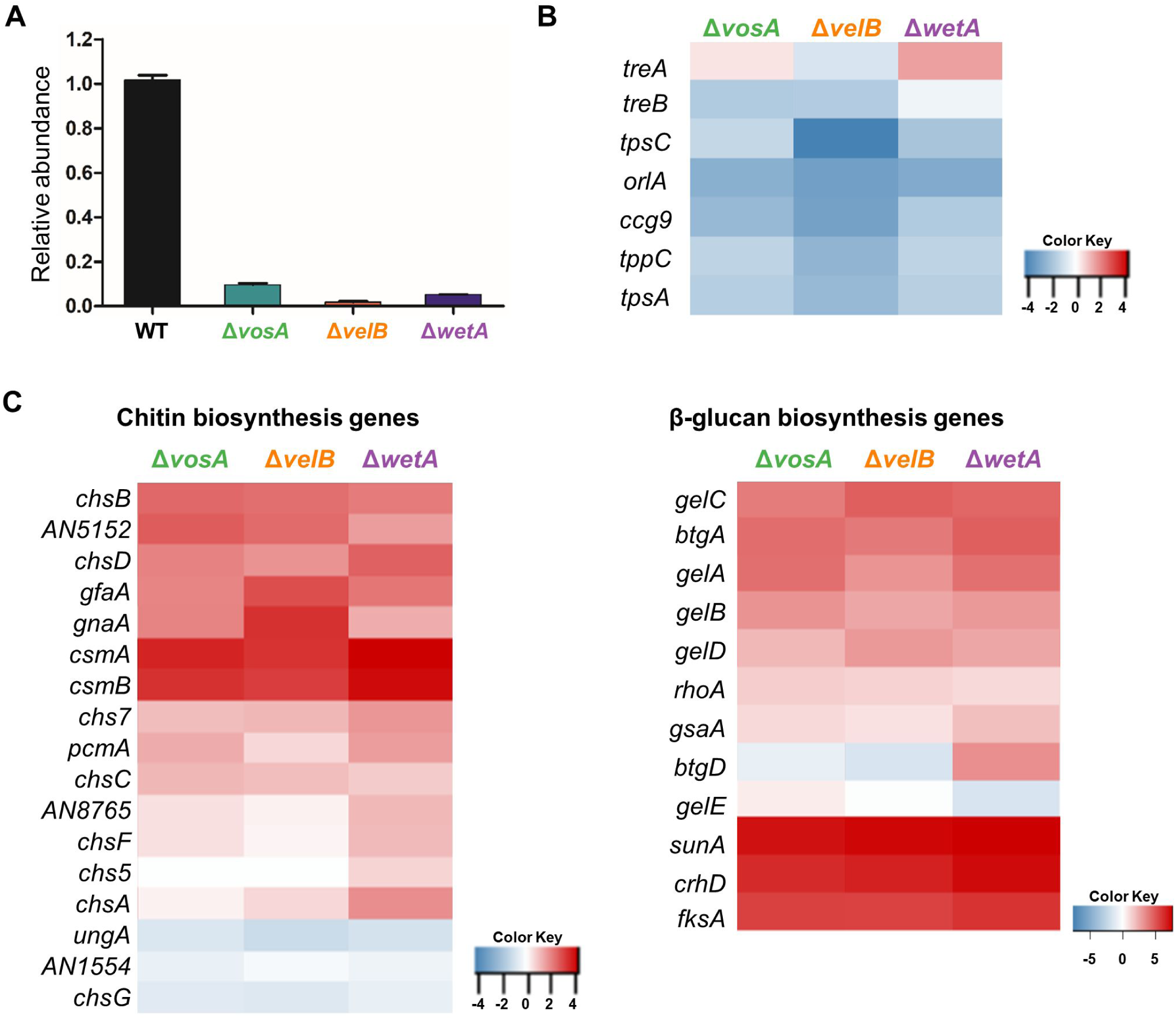
Regulatory effects of VosA, VelB, and WetA on trehalose, chitin, and β-1,3-glucan biosynthesis in *A. nidulans* conidia. **(A)** The amount of conidial trehalose in *A. nidulans*. **(B-D)** Levels of mRNA of the genes associated with trehalose levels (B), chitin biosynthesis (C), and β-1,3-glucan biosynthesis (D) in the Δ*vosA*, Δ*velB*, and Δ*wetA* conidia.

### Alterations to primary metabolites in Δ*vosA*, Δ*velB*, and Δ*wetA* conidia

As mentioned above, the deletion of *vosA, velB*, or *wetA* led to alterations in the mRNA expression of genes involved in metabolic processes (glycerol metabolic process, ketone metabolic process, and amino sugar metabolic process) and amino acid metabolism **(Table S6)**, implying that the amounts of primary metabolites may be affected by the absence of *vosA, velB*, or *wetA* in conidia. To test this hypothesis, the abundances of several primary metabolites involved in the tricarboxylic acid (TCA) cycle and amino acid biosynthesis were examined in WT and mutant conidia **(Figure 4)**. The abundances of pyruvate, α-ketoglutarate, and malate were increased in the conidia of the three null mutants. Acetyl-CoA and succinate were decreased in both Δ*vosA* and Δ*velB*, but not Δ*wetA*, mutant conidia. The amounts of lactate in both Δ*vosA* and Δ*velB* mutant conidia were significantly high, compared with the WT conidia.

**Figure 4.**
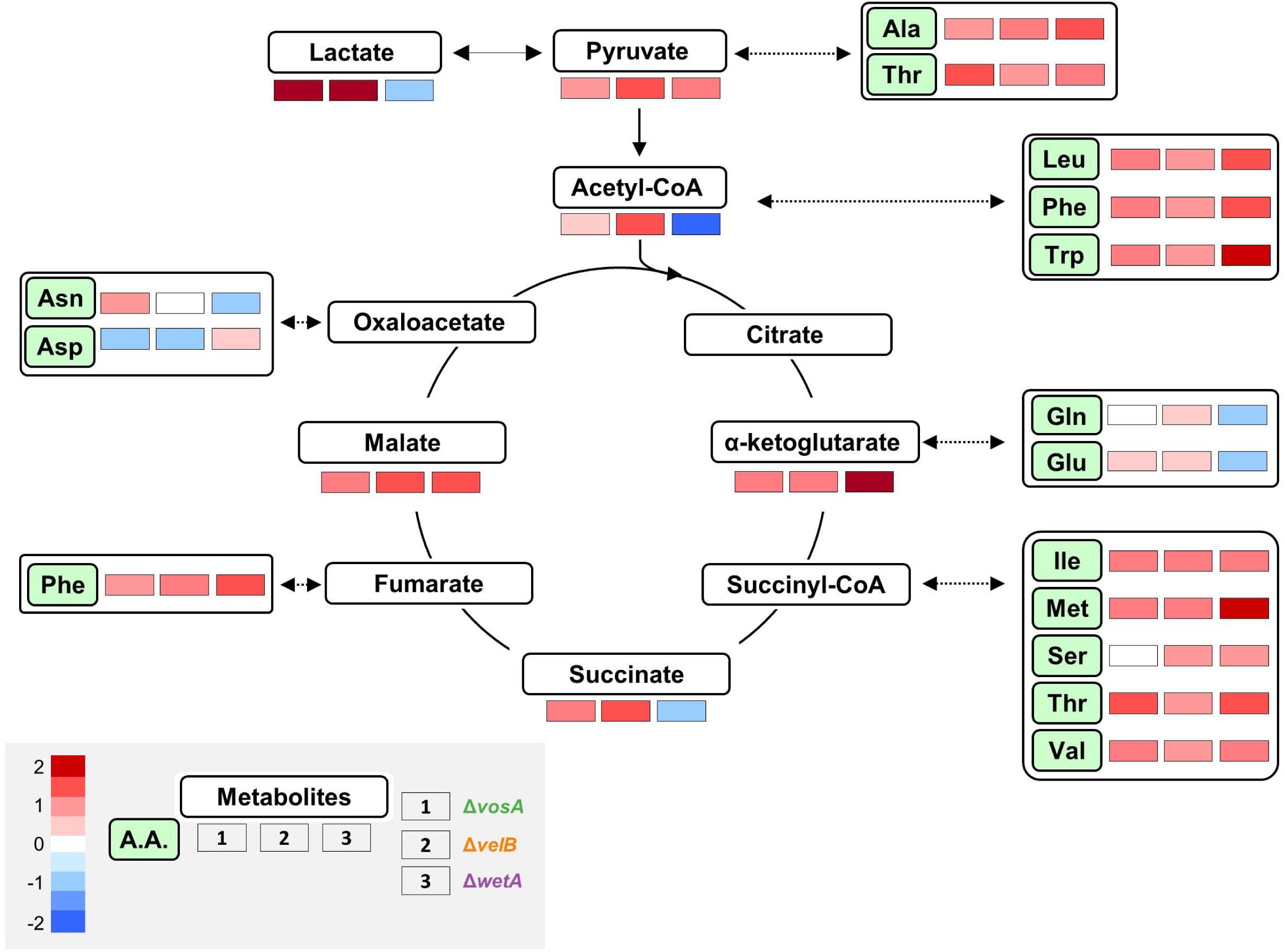
The roles of VosA, VelB, and WetA in primary metabolism of *A. nidulans* conidia. Maps of primary metabolites involved in the TCA cycle and amino acid biosynthesis in WT, Δ*vosA*, Δ*velB*, and Δ*wetA* conidia. Levels of identified primary metabolites produced in the WT and null mutant conidia.

The abundance of 13 amino acids (alanine, isoleucine, methionine, leucine, phenylalanine, tryptophan, valine, threonine, serine, asparagine, glutamine, aspartate, and glutamate) was affected in at least one null mutant. Moreover, levels of nine amino acids were high in all three mutant conidia. The effects of deleting *vosA*/*velB* or *wetA* on the abundance of glutamate, glutamine, aspartate, and asparagine differed. Deletion of *wetA* caused decreased levels of glutamate, glutamine, and asparagine in conidia, whereas, levels of these amino acids were increased or not affected by the absence of *vosA* or *velB*. The genes involved in the biosynthesis of these amino acids and primary metabolites were differentially regulated in the three null mutants. Overall, these results suggest that the regulatory networks of primary metabolites and amino acids are diverse in the three null mutants.

### The abundance of secondary metabolites in Δ*vosA*, Δ*velB*, and Δ*wetA* conidia

Previous studies found that these three transcription factors are important for production of several secondary metabolites in *Aspergillus* species (33, 35, 39). In addition, according to the GO analysis results, deletion of *vosA, velB*, or *wetA* results in alteration of mRNA expression of biosynthetic gene clusters involved in the production of multiple secondary metabolites, including monodictyphenone, sterigmatocystin and asperfuranone **(Figures 1 and S2; see also Table S7)**. To elucidate the conserved and divergent regulatory effects of secondary metabolism in the three conidia mutants, the secondary metabolites were extracted and subjected to liquid chromatography-mass spectrometric (LC/MS) analysis. A principal component analysis showed differences between the four different conidia samples **(Figure S3)**. The secondary metabolite content of the WT conidia was relatively similar to that of the Δ*wetA* conidia, indicating similar abundances and types of secondary metabolites. Conidia from the Δ*vosA* and Δ*velB* mutants clustered far apart, which suggested that a unique set of secondary metabolites or different levels of metabolites were expressed and extracted. This is interesting considering the two TFs can interact and their binding motifs and regulated gene lists were so similar to one another **(Figures 1A and 2A)**.

Next, we applied analysis of variance to identify the most different molecular entities detected as mass/charge (m/z) value and retention time (RT) pairs in the LC/MS analysis-derived metabolomics data. As shown in **Figure 5**, the abundance of several secondary metabolites was different in the positive or negative ionization modes. For example, the abundance of arugosin A was high in the Δ*wetA* conidia, compared with the WT conidia, but not in the Δ*vosA* and Δ*velB* mutant conidia.

**Figure 5.**
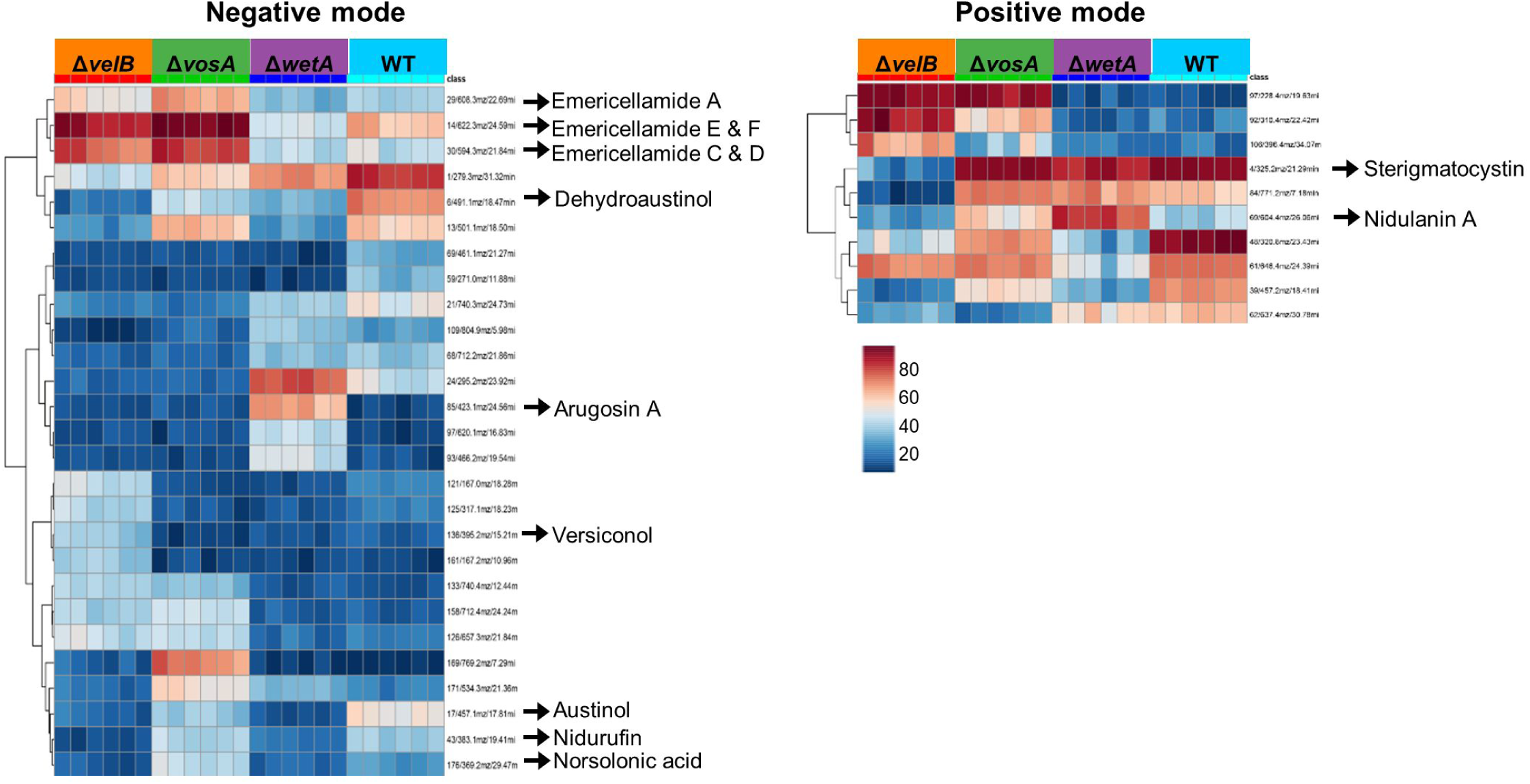
Levels of secondary metabolites in the Δ*vosA*, Δ*velB*, and Δ*wetA* conidia. Differentially regulated secondary metabolites in WT, Δ*vosA*, Δ*velB*, and Δ*wetA* conidia. The heatmap is color-coded and represents high abundance (red) or low abundance (blue) of ions/retention time pairs detected by LC/MS analysis.

To further dissect the roles of VosA, VelB, and WetA in secondary metabolism, we focused on some known secondary metabolites including sterigmatocystin, emericellamide, and austinol **(Figure 6)**. Sterigmatocystin is a precursor of aflatoxins and its biosynthetic gene cluster and intermediates have previously been studied (40, 41). The amount of sterigmatocystin in the Δ*velB* conidia was significantly decreased compared with that in the WT conidia, but the Δ*vosA* and Δ*wetA* conidia contained similar amounts of sterigmatocystin **(Figure 6A)**. However, the amounts of sterigmatocystin intermediates were different in Δ*vosA* and Δ*wetA* conidia. Levels of norsolorinic acid and nidurufin were low in the Δ*velB* and Δ*wetA* conidia, while the amount of versiconol was high only in the Δ*velB* conidia. The RNA-seq results indicated that the mRNA levels of almost all of the genes in the sterigmatocystin gene cluster were increased in both the Δ*vosA* and Δ*wetA* conidia, whereas the mRNA expression of these genes in the Δ*velB* conidia was less consistent. In particular, the mRNA levels of *stcL, stcN, stcQ, stcS, stcT, stcU, stcV*, and *stcW* were decreased in the Δ*velB* conidia, compared with the WT conidia. These results suggest that VosA and VelB play diverse roles in the regulation of sterigmatocystin biosynthesis.

**Figure 6.**
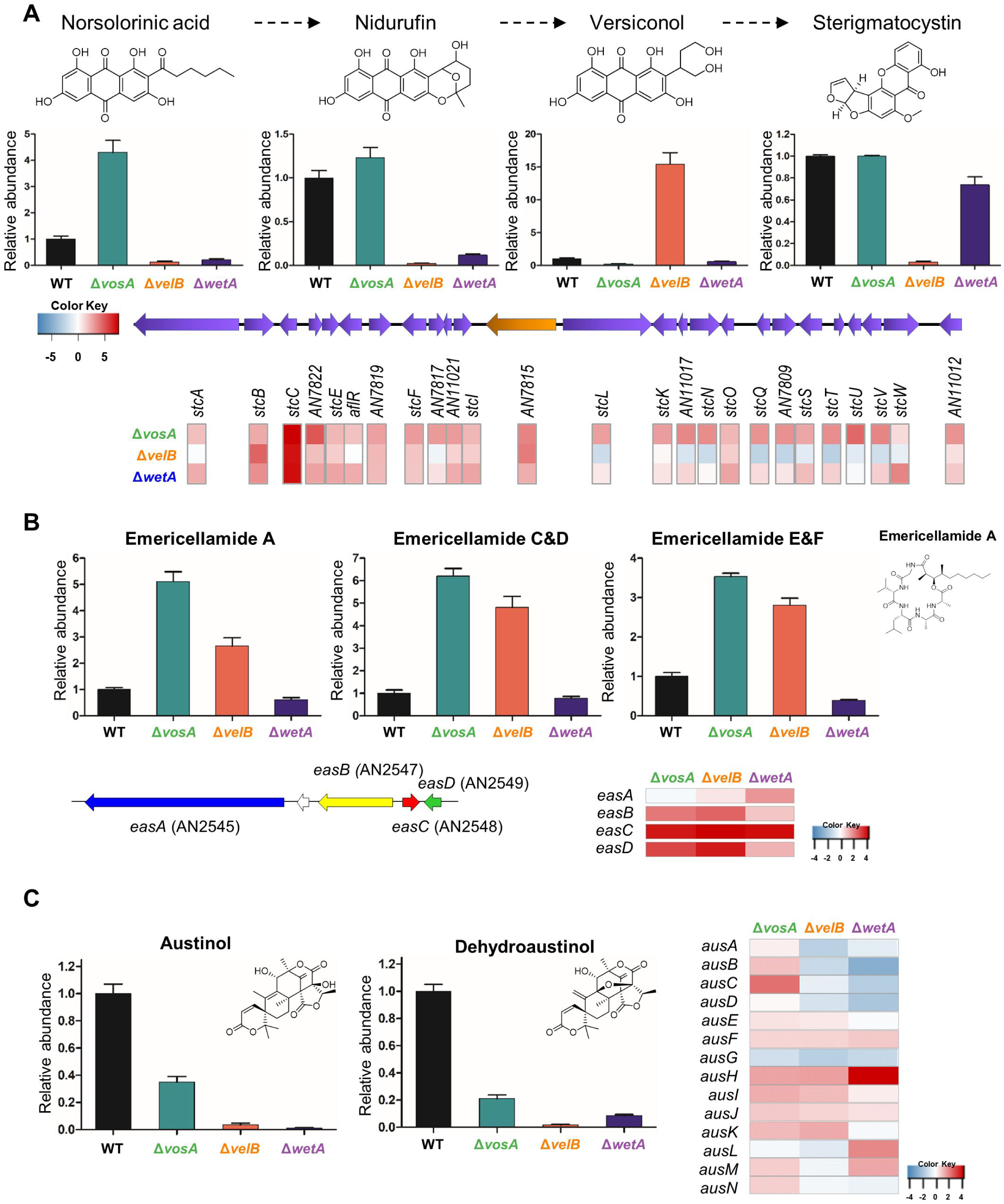
The regulation of key secondary metabolites in Δ*vosA*, Δ*velB*, and Δ*wetA* conidia of *A. nidulans*. **(A)** *Top panel*: The chemical structures of the compounds. *Middle panel*: The abundance of norsolorinic acid, nidurufin, versiconol, and sterigmatocystin in WT, Δ*vosA*, Δ*velB*, and Δ*wetA* conidia. *Bottom panel*: The sterigmatocystin gene cluster and differentially expressed genes involved in sterigmatocystin biosynthesis in Δ*vosA*, Δ*velB*, and Δ*wetA* conidia. **(B)** *Top panel*: The abundance of emericellamide in WT, Δ*vosA*, Δ*velB*, and Δ*wetA* conidia with the emericellamide A structure. *Bottom panel*: The emericellamide gene cluster and mRNA expression of genes associated with emericellamide biosynthesis in Δ*vosA*, Δ*velB*, and Δ*wetA* conidia. **(C)** *Left panel*: The abundance of austinol and dehydroaustinol in WT, Δ*vosA*, Δ*velB*, and Δ*wetA* conidia with their structures. *Right panel*: The austinol gene cluster and mRNA expression of genes associated with austinol biosynthesis in Δ*vosA*, Δ*velB*, and Δ*wetA* conidia.

Emericellamide compounds are cyclopeptides that are produced by several *Aspergillus* species (42, 43). The abundance of these compounds, relative to WT production, was high in Δ*vosA* and Δ*velB* conidia and mRNA levels of *easA-D* were also high in both mutant conidia, implying that VosA and VelB repress emericellamide biosynthesis in WT conidia **(Figure 6B)**. In the Δ*wetA* conidia, however, mRNA expression of the emericellamide gene cluster was increased, but the quantity of emericellamide compounds did not increase, suggesting that the regulatory mechanism of emericellamide biosynthesis in the Δ*wetA* conidia is more complex compared to the influence of Δ*vosA* and Δ*velB* in emericellamide production in conidia. In the three types of null mutant conidia, the abundance of two fungal meroterpenoids, austinol and dehydroaustinol (44), was decreased, compared with the WT conidia **(Figure 6C)**. Furthermore, the expression of several austinol cluster genes was decreased in the Δ*velB* and Δ*wetA* conidia. Taken together, these results demonstrate that the ways in which VosA, VelB, and WetA govern the expression of secondary metabolite gene clusters, and the production of their associated metabolites, in *A. nidulans* conidia are divergent from one another.

## Discussion

Asexual developmental processes in filamentous fungi are regulated by a variety of transcription factors (6). These transcription factors orchestrate the spatial and temporal transcriptional expression of development-specific genes, leading to physiological and metabolic changes. During the processes of conidia formation from phialides and conidial maturation, conidial-specific transcription factors including VosA, VelB, and WetA regulate spore specific gene expression patterns and metabolic changes (25, 30). In this study, we investigated the transcript and metabolite changes that are regulated by VosA, VelB and WetA in *A. nidulans* conidia.

Transcriptomic analyses indicated that about 20% of the *A. nidulans* genome (2143 genes) is differentially expressed in Δ*vosA*, Δ*velB*, and Δ*wetA* mutant conidia. ChIP-seq results identified 66 direct target genes are shared between VosA, VelB, and WetA in conidia. These results offered some explanation of how these transcription factors control phenotypic changes in conidia. First, the deletion of *vosA, velB*, or *wetA* caused increased mRNA expression of certain development-specific genes including *abaA, brlA, flbA, flbC, nsdC, nosA*, and *mpkB*, which are involved in formation of asexual and sexual structures during the early and middle stages of conidia formation, but decreased transcript accumulation of spore-specific genes such as *vadA, catA, wA, conF, conJ, cetA, cetJ*, and *cetL*, which are important for conidial germination, morphogenesis, and dormancy. Another important phenotype of the Δ*vosA*, Δ*velB*, and Δ*wetA* mutant conidia was the differences in conidial wall integrity and the components of the conidial wall (25, 30). As shown **Figure 3**, most of the genes involved in chitin and β-glucan biosynthesis were upregulated in all three mutant conidia. Dynamic expression of these genes is required mainly for the remodeling of the cell wall during isotropic growth and mobilization of energy for differentiation (45), but is not required in dormant conidia. However, by altering the mRNA expression of these genes in the mutant conidia, the dormancy of conidia could be broken, affecting long-term viability, as well as conidial germination.

Another feature of fungal spores is their ability to resist various environmental stresses (1). However, Δ*vosA*, Δ*velB*, and Δ*wetA* mutant conidia are more sensitive to several environmental stresses (25, 34). It is speculated that this is regulated through alterations to the expression of genes involved in environmental stress tolerance. The data we show here support this hypothesis. First, these regulators govern the mRNA expression of genes involved in the trehalose biosynthetic pathway, thereby affecting the amount of conidial trehalose, a key component in stress protection and fungal virulence (46). Second, VosA, VelB, and WetA directly or indirectly regulate genes previously associated with stress responses. CatA is a spore-specific catalase and, compared with WT spores, *catA* deletion mutant spores are sensitive to oxidative stress (47). AtfB is a bZIP transcription factor (48) and the AtfB homolog is crucial to the stress response in *A. oryzae* conidia (49). These two genes are putative direct target genes of the three regulators reported in this study, and the mRNA of *catA* and *atfB* can be positively regulated by VosA, VelB, and WetA in conidia **(Figure 2 and Table S3)**. Along with these genes, the mRNA levels of *hogA*, a key component for osmotic stress signaling (50), was downregulated in all mutant conidia. These results contribute to our understanding of the ways in which these three regulators influence the environmental stress response in conidia.

VosA, VelB, and WetA are key functional regulators in the formation of conidia and control spore-specific gene expression. However, our data has shown that their gene regulation networks are slightly different. RNA-seq results found that VosA and VelB co-regulate the expression of spore-specific genes. Importantly, the predicted VbRE is quite similar to the predicted VoRE **(Figure 2A)**. In addition, biochemical results from previous studies (27, 34) suggested that VosA and VelB form a hetero-complex in asexual spores. However, WetA is not directly related to VosA and VelB. WetA’s putative binding site is different than the VosA/VelB binding site. Moreover, The WetA peak associated genes and the VosA/VelB peak associated genes did not overlap much. These results imply that WetA-mediated gene regulation may be different to the VosA or VelB-mediated gene regulatory network.

During the asexual development of *A. nidulans*, the abundance of amino acids other than phenylalanine changes and the expression of genes related to amino acid biosynthesis is altered (51). Overall, our analyses confirmed that the amount of most amino acids, and the expression of related genes, increased in all mutant spores. In addition, the abundance of metabolites involved in the TCA cycle increased in all mutant conidia. However, the abundance of some primary metabolites such as glutamate, glutamic acid, lactate, and acetyl-CoA was decreased in the Δ*wetA* conidia **(Figure 4)**. It is not yet clear how these metabolic changes affect spore production and maturation, and further studies will be needed to understand this.

An important finding in this study are the mechanisms by which VosA, VelB, and WetA regulate the production of secondary metabolites, especially sterigmatocystin, in conidia. The process of sterigmatocystin production and its regulation involves 25 genes, and this metabolite is produced via steps involving several intermediate products. In Δ*vosA* conidia, the mRNA expression of sterigmatocystin gene clusters was induced and the amount of sterigmatocystin produced were similar to those in the WT conidia. These results were similarly observed in sexual spores (33).While the Δ*vosA* conidia contained sterigmatocystin, the metabolite was not detected in Δ*velB* conidia. We reported that the VosA-VelB complex is a functional unit in conidia, but this particular result indicates that VosA and VelB play different roles in sterigmatocystin production. It is possible that VelB forms another complex, such as the VelB-VeA-LaeA complex(39), to participate in sterigmatocystin production in conidia. In the Δ*velB* conidia, another important finding was that the mRNA expression of genes including *stcB, stcC, stcF*, and *stcI*, which are associated with the early stages of sterigmatocystin biosynthesis, was increased, and the amount of versiconol, a putative sterigmatocystin/aflatoxin intermediate, was also increased, in comparison with the wild type. However, mRNA levels of genes associated with the late phase of sterigmatocystin biosynthesis such as including *stcL, stcN, stcQ*, and *stcT* were decreased in Δ*velB* conidia. It might be possible that VelB (or VelB/VeA/LaeA) can regulate some expression of sterigmatocystin gene clusters by epigenetic means rather than through the canonical method of *aflR* expression or activity.

In conclusion, this study provides a systematic dissection of the gene regulatory network and molecular mechanisms of VosA, VelB, and WetA **(Figure 7)**. In conidia, VosA, VelB, and WetA directly or indirectly control the expression of spore-specific or development-specific genes, thereby altering conidia-wall integrity and conidial viability. In addition, these transcription factors regulate multiple secondary metabolite gene clusters, thus inducing secondary metabolic changes. These results provide an advance in the knowledge of conidial formation and will provide the basis for future insights into spore formation in other filamentous fungi.

**Figure 7.**
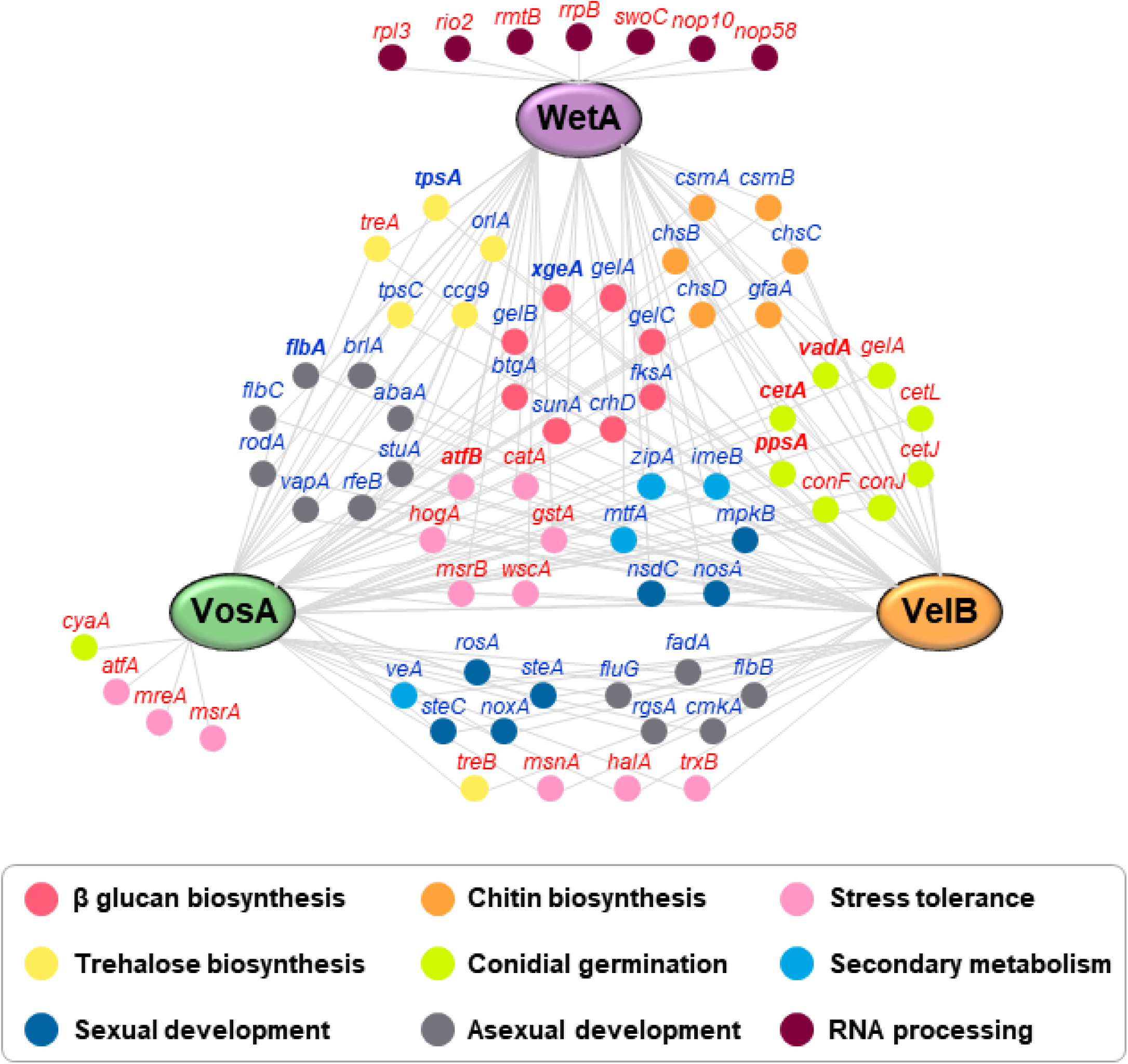
Proposed gene regulatory network of VosA, VelB, and WetA in conidia. The network represents the interactions between VosA/VelB/WetA and their target genes. Gene names in bold typeface are direct target genes of all three TFs. Gene names in red or blue are induced or repressed genes by VosA, VelB, and/or WetA, respectively. in conidia.

## Materials and methods

### Strains, media, and culture conditions

The fungal strains used in this study are listed in **Table 1**. Fungal strains were grown on solid or liquid minimal media with 1% glucose (MMG) and appropriate supplements for general purposes as previously described (52). For conidium samples, WT and mutant conidia were inoculated onto solid MMG plates and incubated for 48 h. Then, conidia were collected from plates using Miracloth (Calbiochem, San Diego, CA, USA) and stored at −80°C.

**Table 1.** *Aspergillus* strains used in this study.

| Strain name | Relevant genotype | References |
| --- | --- | --- |
| FGSC4 | *A. nidulans* wild type, *veA*<sup>+</sup> | FGSC<sup>a</sup> |
| THS15 | *pyrG89; pyroA4; ΔvosA::AfupyrG*<sup>+</sup>; *veA*<sup>+</sup> | (27) |
| THS16 | *pyrG89; pyroA4; ΔvelB::AfupyrG*<sup>+</sup>; *veA*<sup>+</sup> | (27) |
| THS20.1 | *pyrG89; pyroA::velB(p)::velB::FLAG<sub>3x</sub>::pyroA*<sup>b</sup>; *ΔvelB::AfupyrG*<sup>+</sup>; *veA*<sup>+</sup> | (27) |
| THS28.1 | *pyrG89; pyroA::vosA(p)::vosA::FLAG<sub>3x</sub>::pyroA*<sup>b</sup>; *ΔvosA::AfupyrG*<sup>+</sup>; *veA*<sup>+</sup> | (27) |
| TMY4 | *pyrG89; pyroA4; ΔwetA::AfupyrG*<sup>+</sup>; *veA*<sup>+</sup> | (25) |

<sup>a</sup> Fungal Genetic Stock Center<sup>b</sup> The 3/4 *pyroA* marker causes the targeted integration at the *pyroA* locus.

### RNA sequencing (RNA-seq) analysis

To isolate total RNA for RNA-seq analysis, total RNA from WT and mutant conidia was extracted using Trizol Reagent (Invitrogen, USA), according to the manufacturer’s instructions with modifications. To remove DNA contamination from the RNA samples, DNase I (Promega, USA) was added and then RNA was purified using an RNeasy Mini kit (Qiagen, USA). Three technical replicates of each sample were analyzed. RNA sequencing was performed as previously described (33). RNA samples were submitted to the University of Wisconsin Gene Expression Center (Madison, WI, USA) for library preparation and sequencing. A strand-specific library was prepared using an Illumina TruSeq strand-specific RNA sample preparation system. The libraries of all the replicates were sequenced using an Illumina HiSeq 2500 system.

Data analysis of the Δ*vosA* and Δ*velB* RNA-seq experiments used the same analysis pipeline as previously described for the Δ*wetA* RNA-seq analysis (25). Reads were mapped to the *A. nidulans* FGSC4 transcriptome using Tophat2 version 2.1.1 (53) and the parameter “--max-intron-length 4000”. On average, 19.9 million reads per sample mapped to the genome, and the number of reads aligning to each gene was counted with HTseq-Count version 0.9.1 (54). DESeq version 1.14.1 (55) was used to determine significantly differentially expressed genes, and genes were considered regulated if they exhibited an adjusted p-value less than 0.05 and either a log2 fold-change greater than one or less than negative one.

### Chromatin immunoprecipitation sequencing (ChIP-seq) analysis

Samples for ChIP-seq analysis were prepared following methods described previously (29, 30). DNA samples from each strain were extracted using a MAGnify Chromatin Immuno-precipitation System (Invitrogen, USA) according to the manufacturer’s protocol with a modification. Two-day-old conidia from the WT strain, strains containing VosA-FLAG, or VelB-FLAG were cross-linked, washed, homogenized with a Mini-Beadbeater 16 (Biospec, USA), sonicated, and separated with centrifugation. The chromatin extracts were incubated with an anti-FLAG antibody-Dynabead complex. Then, samples were eluted from the beads at 55°C using Proteinase K. Enriched DNA was purified using DNA Purification Magnetic Beads. DNA samples from each strain were submitted to the University of Wisconsin Gene Expression Center (Madison, WI). Libraries were prepared using a TruSeq ChIP library preparation kit (Illumina, CA). The libraries of all the replicates were sequenced using an Illumina HiSeq 2500 system.

Raw reads were trimmed using Trimmomatic version 0.36 (56) and the parameters “ILLUMINACLIP:2:30:10 LEADING:3 TRAILING:3 SLIDINGWINDOW:4:15 MINLEN:36”. Trimmed reads were mapped to the *A. nidulans* A4 genome using version 0.7.15 of BWA-MEM (57) and shorter split hits were marked as secondary alignments. Mapped reads with MAPQ values less than 1, as well as unmapped, secondarily aligned, supplementary, and duplicated reads, were discarded with SAMtools version 1.6 (58). On average, 2.3 million and 7.2 million reads per sample were used for peak calling in the VosA and VelB experiments, respectively. Mapped reads that survived our filter were pooled and extension sizes were estimated with version 1.15.2 of SPP (59, 60). Peaks were called with MACS2 (61) version 2.1.2 using the extension sizes estimated by SPP, a genome size of 2.93e7, and the “--nomodel” parameter. Peaks with a fold-change greater than 2.0 and a q-value less than 0.001 were further analyzed. Peak lists were combined from both of the VosA biological replicates, as >99% of the peaks from the first replicate were found in the second replicate. Motifs were identified in the 100 bp of sequences surrounding each peak summit using MEME-ChIP (62). Motifs that occurred zero or once in the sequences around the peaks and were 4–21 nts in length were further analyzed.

### Functional enrichment analysis

Enriched terms from the “GO Biological Process”, “KEGG”, “InterPro”, and “Pfam” databases were identified using the tools available at AspGD (63), FungiDB (64), and ShinyGO v0.60 (65). Unless otherwise stated, default settings were used in ShinyGO v0.60. The settings were as follows: Database: *Emericella nidulans* STRINGdb, P-value cutoff (FDR): 0.05, # of most significant terms to show: 30.

### Primary metabolite analysis

WT, Δ*wetA*, Δ*vosA*, and Δ*velB* mutant conidia were inoculated onto solid MMG plates and incubated for 48 h, and then fresh conidia were harvested using Miracloth with HPLC-grade water. For each sample, 2 × 10^8^ conidia were mixed with 500 μl HPLC-grade acetonitrile:methanol:water (40:40:20, v/v) and 300 μl beads, homogenized by the Mini Bead beater, and centrifuged. The supernatant was filtered using a 0.45 µm PTFE Mini-UniPrep filter vial (Agilent) and collected and immediately snap frozen with liquid nitrogen. The samples were stored at −80°C until primary metabolite analysis.

The samples were then analyzed as described previously (66, 67). Samples were analyzed using an HPLC-MS system consisting of a Dionex UHPLC coupled by electrospray ionization (ESI; negative mode) to a hybrid quadrupole-high-resolution mass spectrometer (Q Exactive orbitrap, Thermo Scientific) operated in full scan mode. Metabolite peaks were identified by their exact mass and matching retention times to those of pure standards (Sigma-Aldrich).

### Secondary metabolite analysis

The conidia of WT, Δ*wetA*, Δ*vosA*, and Δ*velB* mutant strains were extracted by adding 1.5 ml of a methanol/acetonitrile (2:1) mixture followed by sonication for 60 min. The suspension was then left overnight before centrifugation at 14,000 rpm for 15 mins. The supernatant (1 ml) was removed, filtered, and evaporated to dryness *in vacuo*. Extracts for the metabolomics analysis were normalized to 10 mg/ml in methanol for LC/MS analysis.

Analytical HPLC was performed using an Agilent 1100 HPLC system equipped with a photodiode array detector. The mobile phase consisted of ultra-pure water (A) and acetonitrile (B) with 0.05% formic acid in each solvent. A gradient method from 10% B to 100% B in 35 min at a flow rate of 0.8 ml/min was used. The column (Phenomenex Kinetex C18, 5 μm × 150 mm × 4.6 mm) was re-equilibrated before each injection and the column compartment was maintained at 30°C throughout each run. All samples were filtered through a 0.45 μm nylon filter before LC/MS analysis.

Extracts from the WT and mutant conidia were analyzed in duplicate on an Agilent 1100 series LC/MS platform (68, 69). Negative mode ionization was found to detect most metabolites. The first 5 min of every run was removed due to a large amount of co-eluting, low molecular weight, polar metabolites. Data sets were exported from Agilent’s Chemstation software as .netCDF files and imported into MZmine 2.38 (70). Peak picking was performed with established protocols (71) resulting in 123 marker ions. Briefly, mass detection was centroid with a 5E2 minimum height. Chromatogram building was limited to peaks greater than 0.1 min with 0.05 m/z tolerance and 1E3 minimum height. Data smoothing was performed at a filter width of five. Chromatogram deconvolution utilized a local minimum search with a chromatographic threshold of 95%, minimum relative height of 10%, minimum absolute height of 3E3, minimum ratio of peak to edge of 1, and peak duration range of 0.1–5.0 min. The data was deisotoped with a 1 ppm m/z tolerance before all treatments were aligned and duplicate peaks combined with a tolerance of 0.1 m/z and 3.0 min RT. Peak finder gap filling was performed with 50% intensity tolerance and 0.1 m/z tolerance. Peak lists were exported to Metaboanalyst (72), where missing values were replaced with half the minimum positive value, data were filtered by interquartile range, and log transformation of the data was employed.

## Data availability

All RNA-seq and ChIP-seq data files are available from the NCBI Gene Expression Omnibus database (*wetA* RNA-seq, GSE114143; *vosA* and *velB* RNA-seq, GSE154639; WetA ChIP-seq, GSE114141; VosA and VelB ChIP-seq, GSE154630).

## Acknowledgments

The work at UW-Madison was supported by the National Institute of Food and Agriculture, United States Department of Agriculture, Hatch project 1009695 (MYW and HM), and by the University of Wisconsin-Madison Office of the Vice Chancellor for Research and Graduate Education (OVCRGE) with funding from the Wisconsin Alumni Research Foundation to JHY.

The work by KHH was supported by the Intelligent Synthetic Biology Center of Global Frontier Projects (2015M3A6A8065838) and by Basic Science Research Program through NRF (NRF-2017R1D1A3B06035312) funded by the Korean government.

The work by HSP was supported by the National Research Foundation of Korea (NRF) grant to HSP funded by the Korean government (NRF-2016R1C1B2010945 and NRF-2020R1C1C1004473).

The work by MKL was supported by the KRIBB Research Initiative Program (KGM5232022). The work by GFN, DAA, and SL was supported by the US National Science Foundation grant CH-1808717.

The work by AR and MEM was supported by National Science Foundation grant DEB-1442113 and a Discovery Grant from Vanderbilt University.

## Supplemental Materials

**Table S1**. DEGs related to asexual development in the null mutants’ conidia.

**Table S2**. DEGs related to signal transduction in the null mutants’ conidia.

**Table S3**. VosA, VelB, and WetA peak-associated DEGs.

**Table S4**. Sixty-six DEGs associated with VosA, VelB, and WetA peaks.

**Table S5**. DEGs involved in conidial-wall integrity in the null mutants’ conidia.

**Table S6**. DEGs involved in amino acid metabolism in the null mutants’ conidia.

**Table S7**. DEGs contained in the secondary metabolism gene clusters in the null mutants’ conidia.

**Figure S1. Summary of DEGs in the Δ*vosA*, Δ*velB*, and Δ*wetA* conidia**.

**Figure S2. Gene Ontology (GO) term enrichment analysis of DEGs in the Δ*vosA*, Δ*velB*, and Δ*wetA* conidia**. (A) The top enriched functional categories of the biological process GO terms of DEG in both Δ*vosA* and Δ*velB* conidia (A), or in Δ*wetA* conidia (B).

**Figure S3. Principle component analysis of the metabolic differences between the conidia metabolites from WT, Δ*vosA*, Δ*velB*, and Δ*wetA* strains**.

## References

1. Ebbole DJ. 2010. The Conidium. Cellular and Molecular Biology of Filamentous Fungi:577–590.

2. Wyatt TT, Wosten HA, Dijksterhuis J. 2013. Fungal spores for dispersion in space and time. Adv Appl Microbiol 85:43–91.

3. Park JH, Ryu SH, Lee JY, Kim HJ, Kwak SH, Jung J, Lee J, Sung H, Kim SH. 2019. Airborne fungal spores and invasive aspergillosis in hematologic units in a tertiary hospital during construction: a prospective cohort study. Antimicrob Resist Infect Control 8:88.

4. Latge JP. 1999. *Aspergillus fumigatus* and aspergillosis. Clin Microbiol Rev 12:310–50.

5. Park H-S, Yu J-H. 2012. Genetic control of asexual sporulation in filamentous fungi. Curr Opin Microbiol 15:669–77.

6. Ojeda-Lopez M, Chen W, Eagle CE, Gutierrez G, Jia WL, Swilaiman SS, Huang Z, Park HS, Yu JH, Canovas D, Dyer PS. 2018. Evolution of asexual and sexual reproduction in the aspergilli. Stud Mycol 91:37–59.

7. Casselton L, Zolan M. 2002. The art and design of genetic screens: filamentous fungi. Nat Rev Genet 3:683–97.

8. Martinelli SD. 1994. *Aspergillus nidulans* as an experimental organism. Prog Ind Microbiol 29:33–58.

9. Adams TH, Wieser JK, Yu J-H. 1998. Asexual sporulation in *Aspergillus nidulans*. Microbiol Mol Biol Rev 62:35–54.

10. Etxebeste O, Espeso EA. 2020. *Aspergillus nidulans* in the post-genomic era: a top-model filamentous fungus for the study of signaling and homeostasis mechanisms. Int Microbiol 23:5–22.

11. Park H-S, Yu J-H. 2016. Molecular Biology of Asexual Sporulation in Filamentous Fungi. In: Hoffmeister D. (eds) Biochemistry and Molecular Biology. In D. H (ed), Biochemistry and Molecular Biology The Mycota (A Comprehensive Treatise on Fungi as Experimental Systems for Basic and Applied Research),. Springer, Cham.

12. de Vries RP, Riley R, Wiebenga A, Aguilar-Osorio G, Amillis S, Uchima CA, Anderluh G, Asadollahi M, Askin M, Barry K, Battaglia E, Bayram O, Benocci T, Braus-Stromeyer SA, Caldana C, Canovas D, Cerqueira GC, Chen FS, Chen WP, Choi C, Clum A, dos Santos RAC, Damasio ARD, Diallinas G, Emri T, Fekete E, Flipphi M, Freyberg S, Gallo A, Gournas C, Habgood R, Hainaut M, Harispe ML, Henrissat B, Hilden KS, Hope R, Hossain A, Karabika E, Karaffa L, Karanyi Z, Krasevec N, Kuo A, Kusch H, LaButti K, Lagendijk EL, Lapidus A, Levasseur A, Lindquist E, Lipzen A, Logrieco AF, et al. 2017. Comparative genomics reveals high biological diversity and specific adaptations in the industrially and medically important fungal genus *Aspergillus*. Genome Biology 18.

13. Etxebeste O, Garzia A, Espeso EA, Ugalde U. 2010. *Aspergillus nidulans* asexual development: making the most of cellular modules. Trends Microbiol 18:569–76.

14. Seo JA, Guan Y, Yu JH. 2006. FluG-dependent asexual development in *Aspergillus nidulans* occurs via derepression. Genetics 172:1535–44.

15. Lee MK, Kwon NJ, Lee IS, Jung S, Kim SC, Yu JH. 2016. Negative regulation and developmental competence in *Aspergillus*. Sci Rep 6:28874.

16. Lee MK, Kwon NJ, Choi JM, Lee IS, Jung S, Yu JH. 2014. NsdD is a key repressor of asexual development in *Aspergillus nidulans*. Genetics 197:159–73.

17. Timberlake WE. 1990. Molecular genetics of *Aspergillus* development. Annu Rev Genet 24:5–36.

18. Mirabito PM, Adams TH, Timberlake WE. 1989. Interactions of three sequentially expressed genes control temporal and spatial specificity in *Aspergillus* development. Cell 57:859–68.

19. Adams TH, Boylan MT, Timberlake WE. 1988. *brlA* is necessary and sufficient to direct conidiophore development in *Aspergillus nidulans*. Cell 54:353–62.

20. Adams TH, Deising H, Timberlake WE. 1990. *brlA* requires both zinc fingers to induce development. Mol Cell Biol 10:1815–7.

21. Andrianopoulos A, Timberlake WE. 1991. ATTS, a new and conserved DNA binding domain. Plant Cell 3:747–8.

22. Andrianopoulos A, Timberlake WE. 1994. The *Aspergillus nidulans abaA* gene encodes a transcriptional activator that acts as a genetic switch to control development. Mol Cell Biol 14:2503–15.

23. Sewall TC, Mims CW, Timberlake WE. 1990. *abaA* controls phialide differentiation in *Aspergillus nidulans*. Plant Cell 2:731–9.

24. Sewall TC, Mims CW, Timberlake WE. 1990. Conidium differentiation in *Aspergillus nidulans* wild-type and wet-white (*wetA*) mutant strains. Dev Biol 138:499–508.

25. Wu MY, Mead ME, Lee MK, Ostrem Loss EM, Kim SC, Rokas A, Yu JH. 2018. Systematic Dissection of the Evolutionarily Conserved WetA Developmental Regulator across a Genus of Filamentous Fungi. mBio 9.

26. Marshall MA, Timberlake WE. 1991. *Aspergillus nidulans wetA* activates spore-specific gene expression. Mol Cell Biol 11:55–62.

27. Park HS, Ni M, Jeong KC, Kim YH, Yu JH. 2012. The role, interaction and regulation of the *velvet* regulator VelB in *Aspergillus nidulans*. PLoS One 7:e45935.

28. Ni M, Yu JH. 2007. A novel regulator couples sporogenesis and trehalose biogenesis in *Aspergillus nidulans*. PLoS One 2:e970.

29. Ahmed YL, Gerke J, Park H-S, Bayram O, Neumann P, Ni M, Dickmanns A, Kim SC, Yu J-H, Braus GH, Ficner R. 2013. The *velvet* family of fungal regulators contains a DNA-binding domain structurally similar to NF-kappaB. PLoS Biol 11:e1001750.

30. Park H-S, Yu YM, Lee M-K, Maeng PJ, Kim SC, Yu J-H. 2015. *Velvet*-mediated repression of beta-glucan synthesis in *Aspergillus nidulans* spores. Sci Rep 5:10199.

31. Park H-S, Yu J-H. 2016. Velvet Regulators in *Aspergillus* spp. Microbiol Biotechnol Lett 44:409–419.

32. Bayram O, Braus GH. 2012. Coordination of secondary metabolism and development in fungi: the velvet family of regulatory proteins. FEMS Microbiol Rev 36:1–24.

33. Kim MJ, Lee MK, Pham HQ, Gu MJ, Zhu B, Son SH, Hahn D, Shin JH, Yu JH, Park HS, Han KH. 2020. The velvet Regulator VosA Governs Survival and Secondary Metabolism of Sexual Spores in *Aspergillus nidulans*. Genes (Basel) 11.

34. Sarikaya Bayram O, Bayram O, Valerius O, Park H-S, Irniger S, Gerke J, Ni M, Han KH, Yu J-H, Braus GH. 2010. LaeA control of *velvet* family regulatory proteins for light-dependent development and fungal cell-type specificity. PLoS Genet 6:e1001226.

35. Wu MY, Mead ME, Kim SC, Rokas A, Yu JH. 2017. WetA bridges cellular and chemical development in *Aspergillus flavus*. PLoS One 12:e0179571.

36. Tao L, Yu JH. 2011. AbaA and WetA govern distinct stages of *Aspergillus fumigatus* development. Microbiology 157:313–326.

37. Eom TJ, Moon H, Yu JH, Park HS. 2018. Characterization of the velvet regulators in *Aspergillus flavus*. J Microbiol 56:893–901.

38. Park HS, Bayram O, Braus GH, Kim SC, Yu JH. 2012. Characterization of the velvet regulators in *Aspergillus fumigatus*. Mol Microbiol 86:937–53.

39. Bayram O, Krappmann S, Ni M, Bok JW, Helmstaedt K, Valerius O, Braus-Stromeyer S, Kwon NJ, Keller NP, Yu JH, Braus GH. 2008. VelB/VeA/LaeA complex coordinates light signal with fungal development and secondary metabolism. Science 320:1504–6.

40. Yu J, Chang PK, Ehrlich KC, Cary JW, Bhatnagar D, Cleveland TE, Payne GA, Linz JE, Woloshuk CP, Bennett JW. 2004. Clustered pathway genes in aflatoxin biosynthesis. Appl Environ Microbiol 70:1253–62.

41. Brown DW, Yu JH, Kelkar HS, Fernandes M, Nesbitt TC, Keller NP, Adams TH, Leonard TJ. 1996. Twenty-five coregulated transcripts define a sterigmatocystin gene cluster in *Aspergillus nidulans*. Proc Natl Acad Sci U S A 93:1418–22.

42. Chiang YM, Szewczyk E, Nayak T, Davidson AD, Sanchez JF, Lo HC, Ho WY, Simityan H, Kuo E, Praseuth A, Watanabe K, Oakley BR, Wang CC. 2008. Molecular genetic mining of the *Aspergillus* secondary metabolome: discovery of the emericellamide biosynthetic pathway. Chem Biol 15:527–32.

43. Oh DC, Kauffman CA, Jensen PR, Fenical W. 2007. Induced production of emericellamides A and B from the marine-derived fungus *Emericella* sp. in competing co-culture. J Nat Prod 70:515–20.

44. Lo HC, Entwistle R, Guo CJ, Ahuja M, Szewczyk E, Hung JH, Chiang YM, Oakley BR, Wang CC. 2012. Two separate gene clusters encode the biosynthetic pathway for the meroterpenoids austinol and dehydroaustinol in *Aspergillus nidulans*. J Am Chem Soc 134:4709–20.

45. Baltussen TJH, Zoll J, Verweij PE, Melchers WJG. 2020. Molecular Mechanisms of Conidial Germination in *Aspergillus* spp. Microbiol Mol Biol Rev 84.

46. Thammahong A, Puttikamonkul S, Perfect JR, Brennan RG, Cramer RA. 2017. Central Role of the Trehalose Biosynthesis Pathway in the Pathogenesis of Human Fungal Infections: Opportunities and Challenges for Therapeutic Development. Microbiol Mol Biol Rev 81.

47. Navarro RE, Stringer MA, Hansberg W, Timberlake WE, Aguirre J. 1996. *catA*, a new *Aspergillus nidulans* gene encoding a developmentally regulated catalase. Curr Genet 29:352–9.

48. Lara-Rojas F, Sanchez O, Kawasaki L, Aguirre J. 2011. *Aspergillus nidulans* transcription factor AtfA interacts with the MAPK SakA to regulate general stress responses, development and spore functions. Mol Microbiol 80:436–54.

49. Sakamoto K, Arima TH, Iwashita K, Yamada O, Gomi K, Akita O. 2008. *Aspergillus oryzae atfB* encodes a transcription factor required for stress tolerance in conidia. Fungal Genet Biol 45:922–32.

50. Han KH, Prade RA. 2002. Osmotic stress-coupled maintenance of polar growth in *Aspergillus nidulans*. Mol Microbiol 43:1065–78.

51. Bayram O, Feussner K, Dumkow M, Herrfurth C, Feussner I, Braus GH. 2016. Changes of global gene expression and secondary metabolite accumulation during light-dependent *Aspergillus nidulans* development. Fungal Genet Biol 87:30–53.

52. Kafer E. 1977. Meiotic and mitotic recombination in *Aspergillus* and its chromosomal aberrations. Adv Genet 19:33–131.

53. Kim D, Pertea G, Trapnell C, Pimentel H, Kelley R, Salzberg SL. 2013. TopHat2: accurate alignment of transcriptomes in the presence of insertions, deletions and gene fusions. Genome Biol 14:R36.

54. Anders S, Pyl PT, Huber W. 2015. HTSeq--a Python framework to work with high-throughput sequencing data. Bioinformatics 31:166–9.

55. Love MI, Huber W, Anders S. 2014. Moderated estimation of fold change and dispersion for RNA-seq data with DESeq2. Genome Biol 15:550.

56. Bolger AM, Lohse M, Usadel B. 2014. Trimmomatic: a flexible trimmer for Illumina sequence data. Bioinformatics 30:2114–20.

57. Li H. 2013. Aligning sequence reads, clone sequences and assembly contigs with BWA-MEM. arXiv 1303.3997.

58. Li H, Handsaker B, Wysoker A, Fennell T, Ruan J, Homer N, Marth G, Abecasis G, Durbin R, Genome Project Data Processing S. 2009. The Sequence Alignment/Map format and SAMtools. Bioinformatics 25:2078–9.

59. Landt SG, Marinov GK, Kundaje A, Kheradpour P, Pauli F, Batzoglou S, Bernstein BE, Bickel P, Brown JB, Cayting P, Chen Y, DeSalvo G, Epstein C, Fisher-Aylor KI, Euskirchen G, Gerstein M, Gertz J, Hartemink AJ, Hoffman MM, Iyer VR, Jung YL, Karmakar S, Kellis M, Kharchenko PV, Li Q, Liu T, Liu XS, Ma L, Milosavljevic A, Myers RM, Park PJ, Pazin MJ, Perry MD, Raha D, Reddy TE, Rozowsky J, Shoresh N, Sidow A, Slattery M, Stamatoyannopoulos JA, Tolstorukov MY, White KP, Xi S, Farnham PJ, Lieb JD, Wold BJ, Snyder M. 2012. ChIP-seq guidelines and practices of the ENCODE and modENCODE consortia. Genome Res 22:1813–31.

60. Kharchenko PV, Tolstorukov MY, Park PJ. 2008. Design and analysis of ChIP-seq experiments for DNA-binding proteins. Nat Biotechnol 26:1351–9.

61. Zhang Y, Liu T, Meyer CA, Eeckhoute J, Johnson DS, Bernstein BE, Nusbaum C, Myers RM, Brown M, Li W, Liu XS. 2008. Model-based analysis of ChIP-Seq (MACS). Genome Biol 9:R137.

62. Machanick P, Bailey TL. 2011. MEME-ChIP: motif analysis of large DNA datasets. Bioinformatics 27:1696–7.

63. Arnaud MB, Chibucos MC, Costanzo MC, Crabtree J, Inglis DO, Lotia A, Orvis J, Shah P, Skrzypek MS, Binkley G, Miyasato SR, Wortman JR, Sherlock G. 2010. The *Aspergillus* Genome Database, a curated comparative genomics resource for gene, protein and sequence information for the *Aspergillus* research community. Nucleic Acids Res 38:D420–7.

64. Stajich JE, Harris T, Brunk BP, Brestelli J, Fischer S, Harb OS, Kissinger JC, Li W, Nayak V, Pinney DF, Stoeckert CJ, Jr., Roos DS. 2012. FungiDB: an integrated functional genomics database for fungi. Nucleic Acids Res 40:D675–81.

65. Ge SX, Jung D, Yao R. 2019. ShinyGO: a graphical enrichment tool for animals and plants. Bioinformatics doi: 10.1093/bioinformatics/btz931.

66. Wang PM, Choera T, Wiemann P, Pisithkul T, Amador-Noguez D, Keller NP. 2016. TrpE feedback mutants reveal roadblocks and conduits toward increasing secondary metabolism in *Aspergillus fumigatus*. Fungal Genet Biol 89:102–113.

67. Ostrem Loss EM, Lee MK, Wu MY, Martien J, Chen W, Amador-Noguez D, Jefcoate C, Remucal C, Jung S, Kim SC, Yu JH. 2019. Cytochrome P450 Monooxygenase-Mediated Metabolic Utilization of Benzo[a]Pyrene by *Aspergillus* Species. mBio 10.

68. Adpressa DA, Stalheim KJ, Proteau PJ, Loesgen S. 2017. Unexpected Biotransformation of the HDAC Inhibitor Vorinostat Yields Aniline-Containing Fungal Metabolites. ACS Chem Biol 12:1842–1847.

69. Adpressa DA, Connolly LR, Konkel ZM, Neuhaus GF, Chang XL, Pierce BR, Smith KM, Freitag M, Loesgen S. 2019. A metabolomics-guided approach to discover *Fusarium graminearum* metabolites after removal of a repressive histone modification. Fungal Genet Biol 132:103256.

70. Pluskal T, Castillo S, Villar-Briones A, Oresic M. 2010. MZmine 2: modular framework for processing, visualizing, and analyzing mass spectrometry-based molecular profile data. BMC Bioinformatics 11:395.

71. Abdelmohsen UR, Cheng C, Viegelmann C, Zhang T, Grkovic T, Ahmed S, Quinn RJ, Hentschel U, Edrada-Ebel R. 2014. Dereplication strategies for targeted isolation of new antitrypanosomal actinosporins A and B from a marine sponge associated-Actinokineospora sp. EG49. Mar Drugs 12:1220–44.

72. Xia J, Sinelnikov IV, Han B, Wishart DS. 2015. MetaboAnalyst 3.0--making metabolomics more meaningful. Nucleic Acids Res 43:W251–7.

